# Correlation and Path Coefficient Analyses in Hot Pepper (Capsicum annum L.)

**DOI:** 10.1101/2022.03.30.486344

**Authors:** Bantayehu Bekele, Yohanes Petros, Tamiru Oljira, Mebeaselassie Andargie

## Abstract

To assess the correlation and find out the direct and indirect effect of yield attributing traits on yield, twenty six accessions and four varieties during the off-season period from November 2016 to May 2017 under irrigation. at Dire dawa, Ethiopia. The experiment was conducted using RCBD design with three replications. Plant height, primary branch per plot, days to fifty percent flowering, stem width, number of fruit per plant, fruit length, days to maturity and number of fruit per plant were positive and highly significantly correlated with dry fruit yield per plot at both genotypic and phenotypic correlation. Plant height (0.1081), number of fruit per plant (0.2610), fruit length (0.4293), stem width (0.4059), pedicel length (0.0122), days to maturity (0.0401) and internode length (0.0227) exerted positive direct effect on dry fruit yield per plot at phenotypic level. Genotypic path analysis showed that days to fifty percent flowering (0.0956), stem width (0.5867), fruit length (0.3671), plant height (0.0754), number of fruit per plant (0.2673) and internode length (0.0079)had positive direct effect. The direct effect of these characters on dry fruit yield per plot indicates that improvement on these traits may increase yield.

## 1. Introduction

Chilli (Capsicum annuum L.) originated from tropical and humid zone of Central and Southern America and belongs to the Solanaceae family with diploid chromosome number 2n = 2x = 24. It is a spice, a fruit vegetable widely cultivated in the world and its importance in human food is capital (Dias et al., 2013). Most of the agronomic characters in crop plants are quantitative in nature. Yield is one such character that results due to the actions and interactions of various component characters (Grafius, 1960). It is also widely recognised that genetic architecture of yield can be resolved better by studying its component characters. This enables the plant breeder to breed for high yielding genotypes with desired combinations of traits (Khan and Dar, 2009). Linear correlation between yield and several of its components can present a confusing picture due to inter-relationships between component characters themselves (Khan and Dar, 2009). The objective of this study was to assess the extent of association of traits among themselves and with yield of various Capsicum annuum components among themselves and with yield.

## 2. Materials and Method

### 2.1. Association of characters

#### 2.1.1. Correlation Coefficient

The correlation coefficients among all possible character combinations at phenotypic (rp) and genotypic (rg) level were estimated employing formula Al-Jibouri *et al*. (1958).

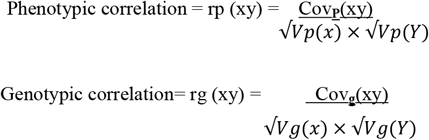

Where,

Covp (xy) and Covg (xy) are phenotypic and genotypic covariance between x and y characters, while Vp (X) and Vg (X) represent variances of X character and Vp (Y) and Vg (Y) denote variances of Y character at phenotypic and genotypic level, respectively.

The coefficient of correlation will be tested for their statistical significance by using t test as,

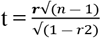

Where n = number of treatment. The calculated value of t was compared with ‘t’ table value at n-2 degree of freedom at 1 and 5 percent level of significance.

#### 2.1.2. Path Coefficient Analysis

Path coefficient analysis was estimated with the formula given by Dewey and Lu (1959).

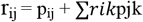

Where:

rij = the association between independent variables (i) and dependent variables (j) as measured by phenotypic and genotypic correlation coefficient.

Pij = component of direct effect of independent variable (i) on the dependent variable (j) as measured by the phenotypic and genotypic path coefficient.

Σ*rikPjk* = is the summation of components of indirect effect of a given independent variable (i) on a given dependent variable (j) via all other independent characters.

The residual effect, which determines how best the causal factors account for the variability of the dependent factor, was calculated as described by Dewey and Lu *et al*. (1959):

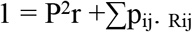

Where, P^2^r is the residual factor, P_ij_ is the direct effect of yield by i^th^ trait on j^th^ trait, and r_ij_ is the correlation of dependent variable with the i^th^ trait.

Small P^2^R value (P^2^R, nearly zero) indicates that the dependent trait considered (yield) is fully explained by the variability in the independent traits.

Higher P^2^R value indicates that some other factors which have not been considered need to be included in the analysis to account fully for the variation in the dependent trait.

## 3. Result and Discussions

### 3.1. Association of Characters for 12 Traits of Chili

Yield of a crop is the result of interaction of a number of interrelated characters. Therefore, selection should be done based on these component characters after assessing their correlation with the yield (Vijaya *et al*. 2014). In the present study, associations of traits for all the 12 quantitative characters were determined for the experimental materials that included 26 accessions and 4 varieties and traits were investigated for their relationship with yield as well as among themselves using genotypic and phenotypic correlation analysis (Table 1 and 2).

**Table 1.** Genotypic correlation coefficient (rg) of yield and yield related 12 quantitative traits of 26 chili accessions and 4 varieties.

| Variables | DF | FS | PB | SW | FG | DM | FL | PL | PH | NFP | IL | DFY |
| --- | --- | --- | --- | --- | --- | --- | --- | --- | --- | --- | --- | --- |
| DF | 1 |  |  |  |  |  |  |  |  |  |  |  |
| FS | -0.404* | 1 |  |  |  |  |  |  |  |  |  |  |
| PB | 0.982** | 0.442* | 1 |  |  |  |  |  |  |  |  |  |
| SW | 0.960** | -0.474 | 0.943** | 1 |  |  |  |  |  |  |  |  |
| FG | 0.329 | -0.467* | 0.321 | 0.373* | 1 |  |  |  |  |  |  |  |
| DM | 0.929** | -0.431* | 0.923** | 0.906** | 0.303 | 1 |  |  |  |  |  |  |
| FL | 0.972** | -0.517* | 0.967** | 0.986** | 0.382* | 0.923** | 1 |  |  |  |  |  |
| PL | 0.193 | -0.226 | 0.133 | 0.172 | 0.257 | 0.024 | 0.138 | 1 |  |  |  |  |
| PH | 0.973** | -0.437* | 0.986** | 0.931** | 0.363* | 0.929** | 0.955** | 0.149 | 1 |  |  |  |
| NFP | 0.970** | -0.490* | 0.979** | 0.943** | 0.349 | 0.929** | 0.966** | 0.137 | 0.987** | 1 |  |  |
| IL | 0.426* | -0.110 | 0.349 | 0.357 | 0.425* | 0.399* | 0.371* | 0.180 | 0.330 | 0.315 | 1 |  |
| DFY | 0.965** | -0.496* | 0.950** | 0.994** | 0.373* | 0.914** | 0.992** | 0.148 | 0.944** | 0.961** | 0.353 | 1 |

\* and \*\* where significant and highly significant at 5% and 1%. DF: Days to fifty percent flowering, FS: Days to first fruit set, PB: Primary branch per plant, SW: Stem width, FG: Fruit girth, DM: Days to maturity, FL: Fruit length, PL: Pedicel length, PH: Plant height, NFP: number of fruit per plant, IL: Internode length, DFY: Dry fruit yield per plot.

**Table 2.**
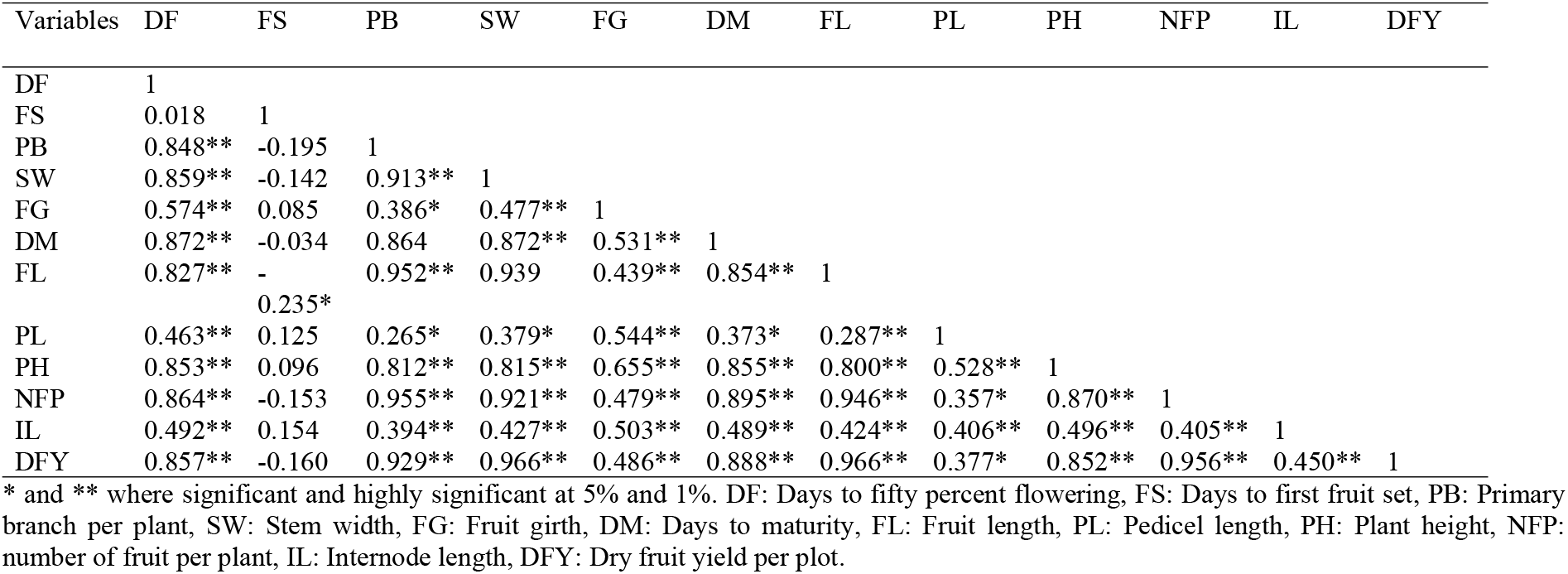
Phenotypic correlation coefficient (rp) of yield and yield related 12 quantitative traits of 26 chili accessions and 4 varieties.

#### 3.1.1. Correlation analysis

##### 3.3.1.1. Genotypic correlation

With genotypic correlations taken as reference, it is found that dry fruit yield per plot is highly significantly and positively correlated with days to fifty percent flowering, primary branch per plant, stem width, days to maturity, fruit length, plant height, and number of fruit per plant. The positive and highly significant correlation indicates a strong association of these traits with dry fruit yield (Demewez *et al*., 2014). Similar findings on chili were reported by (Abraham *et al*., 2017; Luetel *et al*., 2013, Yatung *et al*., 2014) for fruit number per plant, (Singh *et al*., 2014; Sharma *et al*., 2010) for fruit length, (Sharma and Sridevi, 2016) for plant height and fruit length, (Yatung *et al*., 2014, Sharma and Sridevi, 2016) number of fruit per plant, (Abraham *et al*., 2017; Yatung *et al*., 2014) for primary branch per plant. These characters can therefore, be used to the advantage of the breeder for selecting productive genotypes. In contrast, to present study Ajjapplavara (2005) reported negative correlation between fruit yield and primary branch per plant. Similarly, dry fruit yield per plot is correlated significantly and positively with fruit girth but non-significant positive with internode length. Kumari *et al*. (2011) reported similar results for fruit girth. Hence, dry fruit yield per plot can be improved by selecting those traits. Nonetheless, fruit dry weight per plot had non-significant negative correlation with days to first fruiting.

Days to fifty percent flowering, fruit girth, days to maturity, and fruit length showed positive significant association for internode length, whereas non-significant positive association was recorded for primary branch per plant, stem width, pedicel length, plant height, and number of fruit per plant. Similar results were reported by (Sharma *et al*., 2010) for pedicel length, (Singh *et al*., 2009) for primary branch per plant, (Shimeles *et al*., 2016) for days to fifty percent flowering. On the other hand, non-significant negative association was recorded with days to first fruit set.

Highly significant positive correlation of days to fifty percent flowering, primary branch per plant, stem width, days to maturity, fruit length, and plant height with number of fruit/plant revealed that improvement in these traits would enhance number of fruit/plant and ultimately yield. While non-significant positive association was recorded for fruit girth and pedicel length. Such results have been reported by Singh *et al*. (2014), Yatung *et al*. (2014), and Shimeles *et al*. (2016) for primary branch per plant, Sharma *et al*. (2009) for pedicel length. On the other hand, significant negative association was recorded with days to first fruit set.

Highly significant Positive correlation was recorded for plant height with days to fifty percent flowering, primary branch per plant, stem width, days to maturity, and fruit length. Similar results were reported by Singh *et al*. (2009) for primary branch per plant, Shimeles *et al*. (2016) for days to fifty percent flowering and fruit length. Moreover, fruit girth had significant positive association with plant height. On the other hand, negative significant association was recorded with days to first fruit set.

The correlation studies revealed that days to fifty percent flowering, primary branch per plant, stem width, and days to maturity gave highly significant positive association with fruit length. This is in tune with the finding of Shimeles *et al*. (2016) for days to fifty percent flowering. Fruit girth possesses significant positive correlation with fruit length. Nonetheless, fruit length had significant negative correlation with days to first fruit set.

Days to maturity had highly significant positive correlation with days to fifty percent flowering, primary branch per plant, and stem width. However, it had significant negative correlation with days to first fruit set. Fruit girth is correlated significantly and positively with stem width. In contrast, days to first fruit set had significant negative correlation with fruit girth.

The trait, stem width had highly significant positive association with days to first fruiting and primary branch per plant. Nonetheless, it is correlated significantly negatively with days to first fruit set. The association between primary branches per plant with days to fifty percent flowering was significant and negative.

To this end the genotypic correlation profiles indicate that days to fifty percent flowering, primary branch per plant, stem width, days to maturity, fruit length, plant height, number of fruit per plant are important yield components. So, it is suggested that dry fruit yield in chili could be increased by giving weightage to those traits and consequent selection would be rewarding.

##### 3.3.1.2. Phenotypic correlation

Phenotypic coefficients of correlation are presented in Table 2. In the present investigation, dry fruit yield per plot had highly significant (p≤ 0.01) positive correlation with all the traits studied except days to first fruit set and pedicel length. These findings are in consonance with the finding of Sood *et al*. (2009) and Yatung *et al*. (2014) for number of fruit per plant, Singh *et al*. (2014), and Sharma *et al*. (2009) for fruit length, Abraham *et al*. (2017), and Yatung *et al*. (2014) for number of primary branch per plant. This indicates that improvement in these traits would enhance dry fruit yield per plot. Moreover, it is correlated significantly and positively with pedicel length.

Internode length was positive and highly significantly correlated with days to fifty percent flowering, primary branch per plant, stem width, fruit girth, days to maturity, fruit length, pedicel length, plant height, and number of fruit per plant. Number of fruit per plant had highly significant positive relationship with days to fifty percent flowering, primary branch per plant, stem width, fruit girth, fruit length, days to maturity, plant height. Moreover, pedicel length had significant positive correlation with number of fruit per plant. Similar results were reported by Bijalwan and Mishra (2013) for primary branch per plant, (Sing *et al*., 2014) for plant height.

The correlation between Plant height and days to fifty percent flowering, primary branch per plant, stem width, fruit girth, days to maturity, fruit length, and pedicel length were observed to be positive and highly significant. These results were in accordance with the findings of (Sing *et al*., 2014) for primary branch per plant. Pedicel length showed highly significant positive association with days to fifty percent flowering and fruit girth. Besides, it showed significant positive association with primary branch per plant, stem width, days to maturity, and fruit length.

Phenotypic correlation coefficient between fruit length and days to fifty percent flowering, primary branch per plant, stem width, fruit girth, and days to maturity were positive and highly significant. Nonetheless, fruit length exhibit significant negative correlation with days to first fruiting. Days to maturity had highly significant positive association with most of the yield components like days to fifty percent flowering, primary branch per plant, stem width, and fruit girth.

Primary branch per plant, stem width had highly significant positive correlation with fruit girth. Moreover, fruit girth correlated significantly positive with primary branch. Days to fifty percent flowering and primary branch per plant were found to have highly significant positive association with stem width. However, stem width correlated non-significant negative association with days to first fruiting.

Highly significant positive associations were observed between days to fifty percent flowering and primary branch per plant. However, primary branch per plant were found to have negative non-significant association with days to first fruit setting. Days to fifty percent flowering and days to fruit set had non-significant positive association.

From the above discussion of phenotypic correlation it may be concluded that there was highly significant positive correlation of dry fruit yield per plot with days to fifty percent flowering, primary branch per plant, stem width, fruit width, days to maturity, fruit length, plant height, number of fruit per plant, and internode length. So, these traits may be utilized for future breeding programme.

#### 3.3.2. Path coefficient analysis of dry fruit yield Chili with other traits

Knowledge of correlation alone is often misleading as the correlation observed may not be always true. Two characters may show correlation just because they are correlated with a common third one (Khan and Dar, 2009). Under such circumstances, path analysis helps in partitioning of correlation coefficients into direct and indirect effects, permitting a critical examination of the relative importance of each trait (Kumar *et al*., 2013). Correlations between yield and yield components were partitioned into direct and indirect effects to know the particular factor responsible for that correlation (Table 3 and 4).

**Table 3.** Estimates of phenotypic direct effects (bold and diagonal) and indirect effects (off diagonal) of traits via other independent

| Variable | DF | FS | PB | SW | FG | DM | FL | PL | PH | NFP | IL | rp |
| --- | --- | --- | --- | --- | --- | --- | --- | --- | --- | --- | --- | --- |
| DF | **-0.0226** | -0.0002 | -0.1711 | 0.3487 | -0.0223 | 0.0349 | 0.3551 | 0.0057 | 0.0922 | 0.2255 | 0.0112 | 0.857** |
| FS | -0.0004 | **-0.0111** | 0.0394 | -0.0579 | -0.0033 | -0.0013 | -0.1007 | 0.0015 | 0.0104 | -0.0400 | 0.0035 | -0.160 |
| PB | -0.0191 | 0.0022 | **-0.2016** | 0.3706 | -0.0150 | 0.0346 | 0.4087 | 0.0032 | 0.0877 | 0.2493 | 0.0090 | 0.929** |
| SW | -0.0194 | 0.0016 | -0.1841 | **0.4059** | -0.0185 | 0.0349 | 0.4033 | 0.0046 | 0.0880 | 0.2405 | 0.0097 | 0.966** |
| FG | -0.0130 | -0.0009 | -0.0778 | 0.1938 | **-0.0388** | 0.0213 | 0.1885 | 0.0066 | 0.0707 | 0.1252 | 0.0114 | 0.487** |
| DM | -0.0197 | 0.0004 | -0.1743 | 0.3540 | -0.0206 | **0.0401** | 0.3668 | 0.0046 | 0.0924 | 0.2338 | 0.0111 | 0.888** |
| FL | -0.0187 | 0.0026 | -0.1920 | 0.3813 | -0.0170 | 0.0342 | **0.4293** | 0.0035 | 0.0865 | 0.2469 | 0.0096 | 0.966** |
| PL | -0.0104 | -0.0014 | -0.0536 | 0.1542 | -0.0211 | 0.0149 | 0.1233 | **0.0122** | 0.0571 | 0.0934 | 0.0092 | 0.377* |
| PH | -0.0192 | -0.0011 | -0.1637 | 0.3307 | -0.0254 | 0.0343 | 0.3438 | 0.0064 | **0.1081** | 0.2273 | 0.0113 | 0.852** |
| NFP | -0.0195 | 0.0017 | -0.1926 | 0.3740 | -0.0186 | 0.0359 | 0.4061 | 0.0044 | 0.0941 | **0.2610** | 0.0092 | 0.955** |
| IL | -0.0111 | -0.0017 | -0.0795 | 0.1733 | -0.0195 | 0.0196 | 0.1823 | 0.0050 | 0.0537 | 0.1057 | **0.0227** | 0.450** |

**Table 4.**
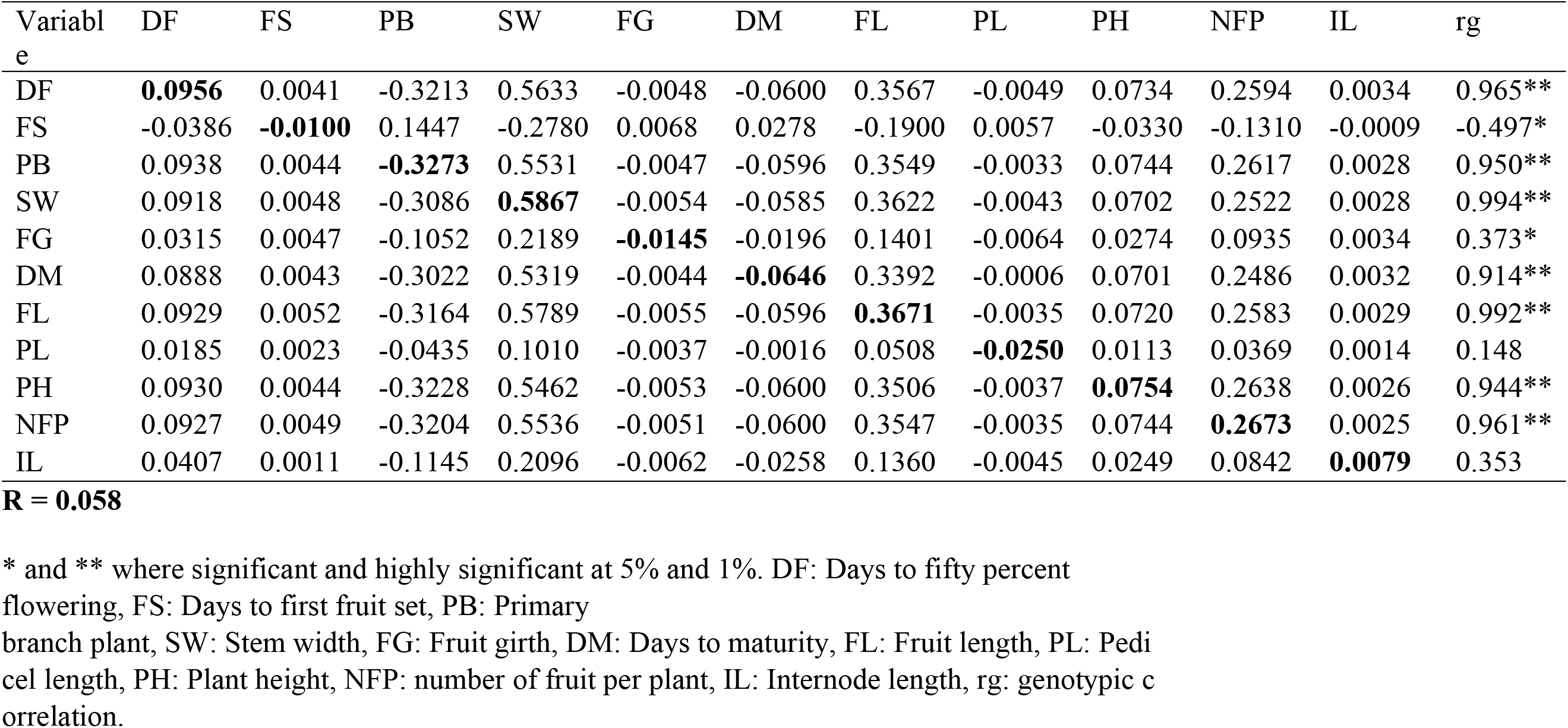
Estimates of genotypic direct effects (bold and diagonal) and indirect effects (off diagonal) of traits via other independent

##### 4.3.2.1. Phenotypic direct and indirect effect of various characters on dry fruit yield per plot

Phenotypic correlations of the different characters were partitioned to path coefficient (Table 3) with the view to identifying important fruit characters having direct effect on yield. The path coefficient analysis at the phenotypic level based on dry fruit yield per plot as dependent variable showed that stem width (0.4059), days to maturity (0.0401), Fruit length (0.4293), pedicel length (0.0122), plant height (0.1081), number of fruit per plant (0.2610), and internode length (0.0227) revealed positive direct effect on dry fruit yield. Thus selection on the basis of those traits would be a paying preposition for evolving high yielding genotypes. On the other hand, days to fifty percent flowering, days to first fruit setting, primary branch per plot, and fruit girth had negative direct effect on dry fruit yield per plot.

Stem width employed direct positive effect (0.4059) on dry fruit yield per plot as well as indirect positive effects via days to first fruit set, days to maturity, fruit length, pedicel length, plant height, number of fruits per plant, and internode length. In contrast to the result of the present study Abraham *et al*. (2017) found negative direct effect of stem width. This contradiction may attributed to the environmental difference of study area. However, stem width exerted negative indirect effect on dry fruit yield per plot through days to fifty percent flowering, primary branch per plant, and fruit girth. Stem width which showed high positive correlation with dry fruit yield per plot is due to its high direct effect and indirect effect through fruit length and number of fruit per plant.

Number of fruits per plant applied positive direct effect (0.2610) and positive indirect effects by means of stem width, days to first fruit set, fruit length, days to maturity, pedicel length, plant height, and internode length. This finding is in harmony with the findings of the previous report by (Abraham *et al*., 2017; Bijalwan and Mishra, 2013; Yatung *et al*., 2014; Sharma *et al*., 2010) for direct effect of number of fruit per plant on yield. The highly significant positive association of number of fruit per plant and dry fruit yield per plot were the result of positive indirect effect of these traits via stem width, fruit length and plant height and also some individual direct effect of number of fruits per plant towards dry fruit yield per plot.

Days to fifty percent flowering had negative direct effect (−0.0226) on dry fruit yield per plot. This trait also exhibited negative indirect effect on dry fruit yield per plot through days to first fruit setting, primary branch per plant and fruit girth. On the other hand, days to fifty percent flowering had an indirect positive effect on dry fruit yield per plot, through stem width, days to maturity, fruit length, pedicel length, plant height, fruit per plant, and internode length.

Days to first fruit setting had negative direct (−0.0194) and indirect effects on dry fruit yield per plot through days to fifty percent flowering, stem width, fruit girth, fruit length, number of fruit per plant. On the other hand, days to first fruit set had an indirect and positive effect on dry fruit yield per plot, through primary branch per plant, days to maturity, pedicel length, plant height, drop per plant, and internode length. Days to first fruit set which correlated non-significant negative with dry fruit yield per plot is due to its negative direct effect as well as negative indirect effect through number of fruit per plant, fruit length, days to maturity, fruit girth, and stem width.

Primary branch per plant had negative direct effect on dry fruit yield per plot. This is in tune with an earlier finding of (Yatung *et al*., 2014). Moreover, it had negative indirect effect via days to fifty percent flowering and fruit girth. In contrast, primary branch per plant expressed positive indirect effect through days to first fruit setting, stem width, days to maturity, fruit length, pedicel length, plant height, number of fruit per plant and internode length. Even though primary branch per plant had negative direct effect its correlation with dry fruit yield per plot is highly significant positive. This is due to high positive indirect effect through stem width, number of fruit per plant, and fruit length.

Phenotypic path coefficient analysis results revealed that fruit girth had negative direct effect and indirect effect via primary branch per plant, days to first fruit setting and days to fifty percent flowering. Supporting evidence of direct negative influence of fruit girth on yield had been reported earlier by (Sharma *et al*., 2010). On the other hand fruit girth exerted positive indirect effect via stem width, days to maturity, fruit length, pedicel length, plant height, number of fruit per plant and internode length. Result obtained from phenotypic correlation indicated that fruit girth had positive highly significant association with dry fruit yield per plot but path coefficient analysis showed that it had negative direct effect on yield. This contradiction is because of positive indirect effect of stem width, fruit length, and number of fruit per plant.

Days to maturity possess positive direct effect on dry fruit yield per plot. Similarly it had positive indirect effect via days to first fruit set, stem width, days to maturity, fruit length, pedicel length, plant height, number of fruit per plant and internode length. On the other hand, it exerted negative indirect effect through fruit girth, primary branch per plant and days to fifty percent flowering. Days to maturity associated highly significantly positive with dry fruit yield per plot. This association is the result of positive direct effect and high indirect effect through stem width, fruit length, and number of fruit per plant.

Fruit length possesses highest positive direct effect on dry fruit yield per plot. These results were in accordance with the findings of (Abraham *et al*., 2017; Bijalwan and Mishra, 2013). Furthermore, it had positive indirect effect via internode length, number of fruit per plant, plant height, pedicel length, days to maturity, stem width and days to first fruit setting. But it exerted negative indirect effect through fruit girth, primary branch per plant and days to fifty percent flowering. The highly significant positive association of fruit length and dry fruit yield per plot were the result of positive indirect effect of these traits via days to maturity, stem width, plant height, and number of fruit per plant and also direct effect of fruit length, towards dry fruit yield per plot.

Pedicel length had positive direct effect on dry fruit yield per plot, as also earlier recorded by Bijalwan and Mishra (2013). Stem width, days to maturity, fruit length, plant height, number of fruit per plant and internode length are traits through which pedicel length exerted indirect positive effect on dry fruit yield per plot. However, pedicel length had negative indirect effect on dry fruit yield per plot via fruit girth, days to first fruit set, primary branch per plant and days to fifty percent flowering.

Plant height had positive direct effect on dry fruit yield per plant. Beside it exerted positive indirect effect via stem width, days to maturity, fruit length, pedicel length, number of fruit per plant and internode length. In contrary, it had negative indirect effect through fruit girth, primary branch per plant, days to first fruit set and days to fifty percent flowering. The result of phenotypic correlation showed that plant height and dry fruit yield per plot had highly significant positive correlation

The number of fruits per plant exhibited direct positive effect and indirect effect through other characters like days to first fruit setting, stem width, days to maturity, fruit length, pedicel length, plant height and internode length. These findings are in consonance with (Sharma *et al*., 2014) for positive direct effect of Number of fruit per plant. Nonetheless, it had negative indirect effect, via fruit girth, days to fifty percent flowering and primary branch per plant. Number of fruit per plant had highly significant positive correlation with dry fruit yield per plot. Path analysis revealed that this association is due to positive direct effect of number of fruit per plant and positive indirect effect through stem width, fruit length, and plant height.

Internode length exerted positive direct effect on dry fruit yield per plot and exerted positive indirect effect through stem width, days to maturity, fruit length, pedicel length, plant height and number of fruit per plant. On the other hand, internode length exerted negative indirect effect through days to first fruit set, primary branch per plant and days to fifty percent fruiting. The residual (0.158) indicated that the independent variables included in this study explained (84.2%) of the total variation in the dependent variables that is dry fruit yield per plot. The unexplained variation in the phenotypic path coefficient was 0.158. It predicted that 84.2% variation at phenotypic level had been determined and further indicated that some more factors need to be considered in this study contributed to dry fruit yield per plot. Thus, few more traits may be considered while selecting the genotypes for high yield.

Overall, the phenotypic path analysis confined that direct effect of stem width, fruit length, number of fruit per plant, plant height, days to maturity, internode length, and pedicel length and negative direct effect of days to fifty percent flowering, days to first fruit set, number of primary branch per plant, and fruit girth should be considered simultaneously for amenability in dry fruit yield of chili selections.

##### 3.3.2.2. Genotypic direct and indirect effect of various characters on dry fruit yield per plot

The path analysis at the phenotypic level may not provide a true picture of direct and indirect causes (Sood *et al*., 2009), and it is advisable to understand the contribution of different traits toward the dry fruit yield/plot at the genotypic level. For path analysis at the genotypic level, dry fruit yield/plot was the dependent variable to all other traits used for correlation and considered as casual variables.

The Genotypic direct and indirect effect of different characters on dry fruit pepper yield is presented in Table 4. The maximum positive genotypic direct effect on dry fruit yield per plot was observed in stem width (0.5867) followed by fruit length (0.3671), number of fruit per plant (0.2673), days to fifty percent flowering (0.0956), plant height (0.0754), inter node length (0.0026). Negative direct effects were recorded for days to first fruit set (−0.01), primary branch per plant (−0.3273), fruit girth (−0.0145), days to maturity (−0.0646), and pedicel length (−0.0250).

Days to fifty percent flowering (0.0956) had direct positive effect and indirect positive effect through days to first fruit set, stem width, fruit length, plant height, number of fruit per plant and internode length. These results were in accordance with the findings of (Bijalwan and Mishra, 2013 and Shimeles *et al*., 2016) who reported positive direct effect of days to fifty percent flowering. On the other hand, it had indirect negative effect through primary branch per plant, fruit girth, days to maturity and pedicel length.

Days to first fruit set (−0.01) possess negative direct effect as well as it exerted negative indirect through days to first fruit set, internode length, number of fruit per plant, days to fifty percent flowering, stem width, stem width and plant height. On the other hand days to first fruit set contribute indirect positively through primary branch per plant, fruit girth, days to maturity and pedicel length to dry fruit yield per plot. Days to first fruit set exhibited negative correlation with yield, which is due to negative direct effect as well as negative indirect effect through days to first fruit set, stem width, fruit length, plant height, number of fruit per plant, and internode length.

Primary branch per plant (−0.3273) had negative direct effect and recorded a high positive indirect effect *via* stem width followed by fruit length, number of fruit per plant, days to fifty percent flowering, plant height, days to first fruit set and internode length. It causes negative indirect effect through fruit girth, days to maturity and pedicel length. Similar results were reported by (Bijalwan and Mishra, 2013; Singh, 2014 and Yatung *et al*., 2014) for negative direct effect of primary branch per plant. Genotypic correlation analysis indicated primary branch per plant as an important character influencing dry fruit yield. However, path coefficient analysis suggested that primary branch per plant had negative direct effect on yield but had indirect positive influence through stem width, fruit length and number of fruit per plant. The apparent contraction is due to the total correlation measures mutual association without causation whereas path coefficient analysis specifies the causes and measures their relative importance.

Stem width which possess highest correlation (0.994) also exhibit highest direct effect (0.5867) on dry fruit yield per plot. Hence, it would be rewarding to lay emphasis on stem width while developing selection strategies towards high yielding varieties. The indirect contribution of stem width was positive *via* fruit length, number of fruit per plant, days to fifty percent flowering, plant height, days to first fruit set and internode length. In contrast, negative indirect effect was observed by stem width through pedicel length, fruit girth, primary branch per plant and stem width. The highest correlation exhibited by stem width is due to highest direct and indirect effect through fruit length and number of fruit per plant.

Fruit girth exhibited (−0.0145) negative direct effect. Such results have been reported by Sharma *et al*. (2010). The indirect contribution of fruit girth was positive and high via stem width and followed by fruit length, number of fruit per plant, days to fifty percent flowering, plant height, days to first fruit set and internode length.

Concerning days to maturity (−0.0646), it had negative direct effect on dry fruit yield per plot. Such results also were reported by (Singh *et al*., 2014). Its indirect effects through stem width, fruit length, number of fruit per plant, days to fifty percent flowering, plant height, days to first fruit set and internode length were positive. But it exerts negative indirect effect only through primary branch per plant.

The direct effect of fruit length on dry fruit yield per plot was (0.3671) positive. Such results have been reported by (Abraham *et al*., 2017; Bijalwan and Mishra, 2013; Yatung *et al*., 2014; Shimeles *et al*., 2016 and Sing *et al*., 2014). It had highest positive indirect effect on dry fruit yield per plot through stem width followed by number of fruit per plant, days to fifty percent flowering, plant height, days to fifty percent flowering and internode length. Also had negative indirect effect on dry fruit yield per plot through primary branch per plant followed by days to maturity, fruit girth and pedicel length. Fruit length and dry fruit yield per plot had highly significant positive correlation which is due to high positive direct effect and indirect effect through stem width and number of fruit per plant as revealed by genotypic path analysis.

Pedicel length had negative direct effect (−0.0250) on dry fruit yield per plot and in direct effect via primary branch per plant and fruit girth. Bijalwan and Mishra (2013 and Sharma *et al*. (2009) reported negative direct effect of pedicel length. On the other way it had positive indirect effect on dry fruit yield per plot through stem width followed by fruit length, number of fruit per plant, days to fifty percent flowering, plant height, days to first fruit set and internode length.

Plant height had positive (0.0754) direct effect on dry fruit yield per plot and indirect positive effect via stem width followed by fruit length, number of fruit per plant, days to fifty percent flowering, days to first fruit set and internode length. Corroborating the findings of present investigation positive direct effect of plant height on yield has also been reported by (Abraham *et al*., 2017; Bijalwan and Mishra, 2013). Nonetheless, plant height exerted negative indirect effect through primary branch per plant followed by days to maturity and fruit girth.

Number of fruit per plant exerted a positive (0.2673) direct effect. This finding has earlier been supported by (Abraham *et al*., 2017; Bijalwan and Mishra, 2013; Yatung *et al*., 2014; Sharma *et al*., 2009; Shimeles *et al*., 2016; and Sing *et al*., 2014). It had positive highest indirect effect through stem width followed by fruit length, days to fifty percent flowering, plant height, days to first fruit set and internode length. In contrast, it exerted negative indirect effect through primary branch per plant followed by days to maturity, fruit girth and pedicel length. Genotypic correlation showed highly significant positive correlation of number of fruit per plant with dry fruit yield per plot. Path analysis revealed that this association is due to positive direct and high positive indirect effect through stem width and fruit length.

Internode length possesses positive (0.0079) direct effect on dry fruit yield per plot. It had positive indirect effect through stem width followed by fruit length, number of fruit per plant, days to fifty percent flowering, plant height and days to first fruit set. In contrast, internode length exerted negative indirect effect via primary branch per plant followed by days to maturity, fruit girth and pedicel length.

Residual effect genotypic (0.058) level indicated that the traits included in the present investigation accounted for most of the variation (94.2%) present in the dependent variable that is dry fruit yield per plot.

In general, from genotypic path analysis, the number of fruits per plant, stem width, fruit length days to fifty percent flowering, plant height, and internode length exhibited the positive direct effect in which one can improve the dry fruit yield through direct selection of either of these characters during yield improvement program. Moreover, stem width score highest positive direct and indirect effect as a result, it should be given more emphasis in the selection aimed at improving dry fruit yield in chili.

## 4. Conclusion

On the basis of correlation studies at both genotypic and phenotypic level and their coefficient of determination, the selection for Stem width, days to fifty percent flowering, primary branch per plant, days to maturity, fruit length, plant height, and number of fruit per plant will be effective for isolating plants with higher yield in chili.

In view at the direct and indirect contributions of component traits towards fruit yield at genotypic and phenotypic path analysis, selection on the basis of yield related traits viz., stem width, fruit length, plant height, number of fruit per plant, and internode length would be a paying preposition in the accessions and varieties included in the study.

